# Integrated systems biology and imaging of the smallest free-living eukaryote *Ostreococcus tauri*

**DOI:** 10.1101/293704

**Authors:** Chuck R. Smallwood, Jian-Hua Chen, Neeraj Kumar, William Chrisler, Samuel O. Purvine, Jennifer E. Kyle, Carrie D. Nicora, Rosanne Boudreau, Axel Ekman, Kim K. Hixson, Ronald J. Moore, Gerry McDermott, William R. Cannon, James E. Evans

## Abstract

*Ostreococcus tauri* is an ancient phototrophic microalgae that possesses favorable genetic and cellular characteristics for reductionist studies probing biosystem design and dynamics. Here multimodal bioimaging and multi-omics techniques were combined to interrogate *O. tauri* cellular changes in response to variations in bioavailable nitrogen and carbon ratios. Confocal microscopy, stimulated Raman scattering, and cryo-soft x-ray tomography revealed whole cell ultrastructural dynamics and composition while proteomic and lipidomic profiling captured changes at the molecular and macromolecular scale.

Despite several energy dense long-chain triacylglycerol lipids showing more than 40-fold higher abundance under N deprivation, only a few proteins directly associated with lipid biogenesis showed significant expression changes. However, the entire pathway for starch granule biosynthesis was highly upregulated suggesting much of the cellular energy is preferentially directed towards starch over lipid accumulation. Additionally, three of the five most downregulated and five of the ten most upregulated proteins during severe nitrogen depletion were unnamed protein products that warrant additional biochemical analysis and functional annotation to control carbon transformation dynamics in this smallest eukaryote.

## Introduction

Microalgae are ubiquitous in oceans and maintain a major carbon sink in the complex world-wide ecosystem. Spanning more than one-billion years of evolution, microalgae are phylogenetically diverse and exhibit varying cellular phenotypes that naturally produce high value metabolites, proteins, carbohydrates and energy dense lipids that can be exploited for a wide array of industrial applications ^1–3^. Due to their high photosynthetic efficiency for energy conversion, and minimal growth requirements consisting of sustainable resources such as marine or brackish media, light, CO_2_ and trace vitamins, microalgae are prime bioproduction platforms ^4,5^.

Triacylglycerol (TAG) lipids are significantly enhanced when microalgae are subjected to cellular stressors such as light, temperature, and nutrient deprivation ^6^. TAG lipids possess nonpolar character and are stored in anhydrous, high-density organelles called lipid bodies, which are desirable for industrial lipid feedstock applications ^7^. For other oleaginous algae, TAG production can be triggered by nutrient deprivation of iron, sulfur, nitrogen, phosphate, or silicon ^6^. In most eukaryotes, combinatorial reduction of these nutrients results in altered levels of growth-associated structural lipids (phospholipids) and energy storage lipids (TAG) products ^8,9^. Unfortunately, in most cases starvation or deprivation can be detrimental to cell viability and overall growth capacity thereby limiting cell biomass yields needed for viable lipid feedstock industrial applications ^10^. While several reports have shown that supplementing additional C sources when combined with N depletion for certain organisms can yield increased growth and lipid accumulation rates compared to strict starvation^2,11,12,13^, a detailed understanding of the interplay between carbon and nitrogen bioavailability is needed to advance bioproduction applications.

Studying phototrophic metabolism in primitive species, such as the prasinophyte *Ostreococcus tauri* can provide convenient opportunities to define minimal and critical metabolic pathways for C transformation^14,15,16^. *O. tauri* is the smallest known free-living eukaryote (~0.8*μ*m in thickness), lacks a cell wall, and thrives in varying photic, toxic and thermal ecosystems^17–19^. It also has a highly condensed genome with only ~8,000 genes^20^ so most reactions are governed by a single enzyme (rather than multiple duplicating function enzymes) which simplifies engineering requirements and interpretation. For example, the canonical model green alga *Chlamydomonas* has 4 copies of Acetyl-CoA Carboxylase (E.C. 6.4.1.2) whereas *Ostreococcus* has only 1 enzyme of that class. In addition, a recent study^21^ reported the genetic diversity associated with large phenotypic differences between *Ostreococcus* strains highlighting the uncharted abundance of genetic biodiversity. These characteristics make *O. tauri* a potential candidate for future industrial applications. However, considerable biodesign efforts will likely be needed to develop an efficient cell factory for controlled and cost-effective bioproduction of lipid feedstocks or other high value metabolites. Here, an integrated analysis of bioimaging, proteomic, and lipidomic characterization was used to investigate *O. tauri* cellular response to varying C:N ratios. Our results provide additional understanding of C storage and energy transformation pathways within the microalgae *O. tauri* and identify new proteins to target for future engineering efforts.

## Results

Other than the bioavailable C:N ratio, all other experimental parameters (e.g., temperature, duration, diurnal cycling, light fluence, etc.) were kept constant throughout this study. The four different media conditions used herein are designated as K6CN, K2CN, K2C and K6C which are listed in decreasing C:N ratio. The normal growth media for *O. tauri* is Keller Media (referenced herein as K2CN media) and contains ~2.5mM bicarbonate and 0.9mM total nitrogen. To streamline interpretation, bicarbonate was kept as the sole media carbon source. For K6CN media the nitrogen content was kept equivalent, but the bicarbonate increased to 6mM providing elevated carbon but normal nitrogen conditions. The use of 6mM bicarbonate as the elevated carbon set point was chosen following an initial growth screen of *O. tauri* in K media supplemented with various levels of bicarbonate from 0-10mM where 6mM showed highest growth response with healthy chlorophyll ratios per cell. The final two media conditions K2C and K6C share the same composition as K2CN and K6CN but are depleted of all nitrogen sources. Thus all subsequent experiments consisted of 4 conditions of decreasing C:N bioavailable ratios from elevated carbon with normal nitrogen (K6CN), to normal carbon with normal nitrogen (K2CN), to normal carbon with no nitrogen (K2C), and elevated carbon with no nitrogen (K6C).

Diurnal cycling with 12:12 hour light:dark cycles was used to synchronize cellular division and growth, and samples were harvested 3 hours after light to dark transition. Cell growth was monitored by absorbance at 750nm (measure of particulates) and 680nm (measure of chlorophyll a) for up to 144 hours (Fig. 1B). Decreases in absorbance at 680nm were typical of K2C and K6C cell cultures relative to K2CN and K6CN. Absorbance at 750nm exhibited similar decreases for N deprived cultures versus N replete. However, when comparing normal C to excess C for either N replete or N deprived conditions, cultures with excess C consistently displayed higher A680 and A750 values. Confocal fluorescence microscopy was used to compare phenotypes for a couple dozen cells from each condition whereas fluorescence activated cell sorting (FACS) allowed quantitative analysis of larger population dynamics every 24 hours. In both case, dramatic increases of neutral lipid (NL) content were detected for K2C and K6C conditions (Fig. 1 C–F). Interestingly K6CN (Fig. 1C) cell cultures only exhibited subtle differences between NL and some increases in phospholipid (PL) intensity indicating some photosynthetic lipid metabolism difference to K2CN (Fig. 1D). Since confocal microscopy can only track dynamics for fluorescently labelled components, label-free cell ultrastructure and composition changes were also evaluated.

**Figure 1.**
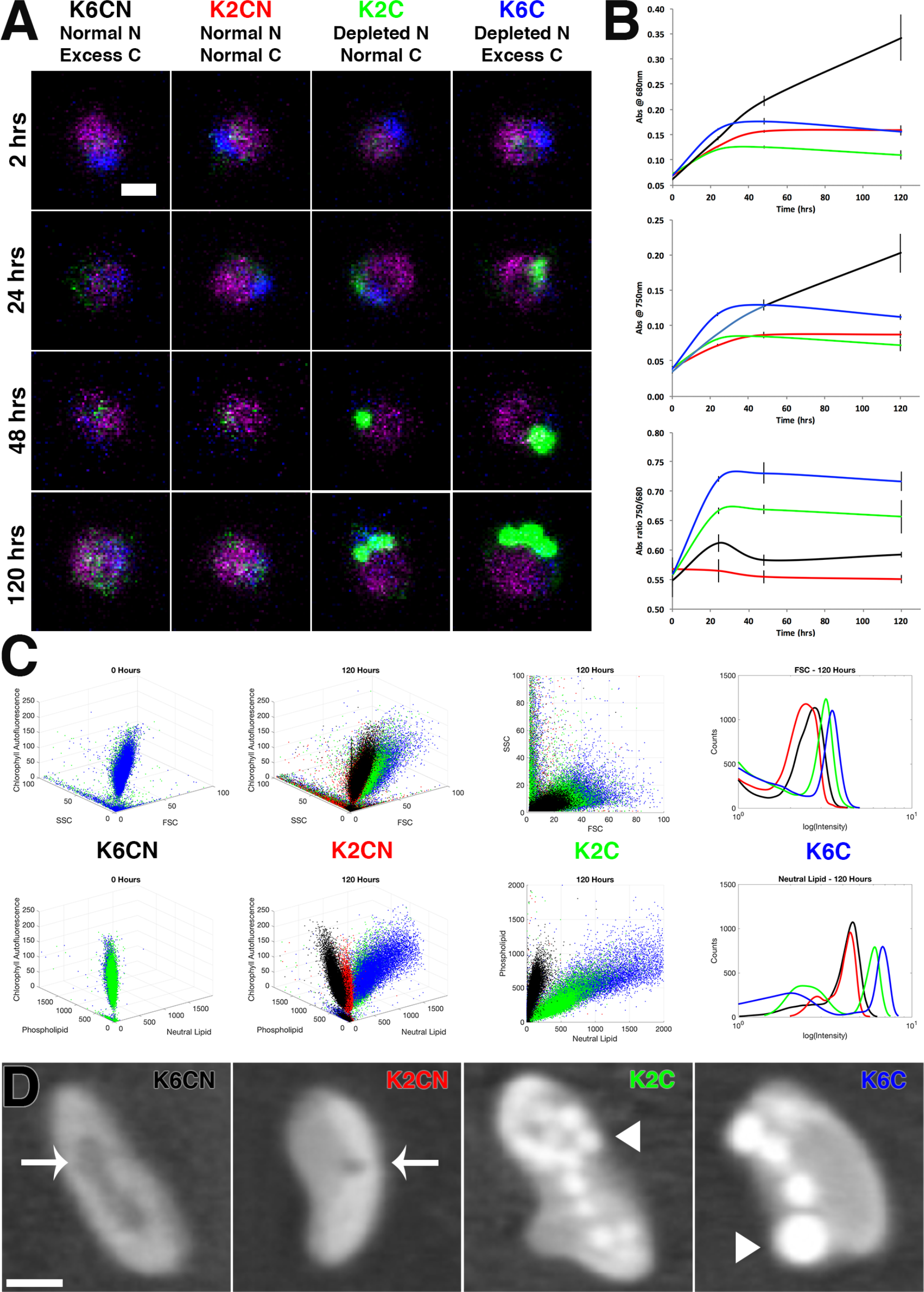
Single cell and population imaging and growth under varying C:N ratios. (A) Confocal microscopy of K6CN, K2CN, K2C, and K6C cultures to observe chlorophyll autofluorescence (magenta), nucleic acid fluorescent staining (blue), neutral lipid fluorescent staining (green) over time. Scale bar represents 1*μ*m scale. (B) Corresponding growth plots for the same cultures monitored over time (error bars represent a replicate of 5) at 750nm and 680nm, and the ratio of the 750/680 as a measure of photosynthetic efficiency. (C) FACS time course study of same cultures comparing SSC (side-scattering) versus FSC (forward scattering) and chlorophyll autofluorescence in 3D plots at 0 hours and 120 hours, then 2D plot of SSC versus FSC at 120 hours, and histogram of FSC at 120 hours. (D) FACS time course study of same cultures comparing phospholipid versus neutral lipid and chlorophyll autofluorescence in 3D plots at 0 hours and 120 hours, then 2D plot of phospholipid versus neutral lipid at 120 hours, and histogram of neutral lipid at 120 hours. (E) Label-free cryogenic soft x-ray nanotomography images of cells cryogenically frozen in microcapillaries. Central slices (1*μ*m scale bar) display intracellular structures common to each culture condition. Chloroplasts and cytoplasm seen in all images. Vacuoles in K6CN and K2CN appear as dark objects indicated with white arrow. Lipid droplets appear as bright white circular objects indicated by white arrowhead.

Cryogenic soft x-ray nanotomography (CSXT) was performed on 72-hour cell cultures to increase resolution and highlight native cellular features of intracellular lipid accumulation (Fig. 1E). Dark subcellular features, assigned as vacuoles due to their linear absorption coefficient (LAC), were observed in the intracellular cytosolic space of cells in K6CN and K2CN culture conditions, but no vacuoles were observed for K2C and K6C cultured cells. The vacuoles were much larger in K6CN compared to K2CN conditions. Based on studies conducted in other algae and plants starch is likely present in the chloroplast^20^. However, intracellular starch was undetected in *O. tauri* cells via X-ray nanotomography likely due to the density appearing similar to or masked by the surrounding tissues with a similar LAC - possibly due to its small cell size or chloroplast packing density. Therefore, stimulated Raman scattering (SRS) microscopy was used as a second label-free imaging method to identify relative abundance of intracellular starch accumulation. Interestingly, starch was detected in the chloroplast for all tested conditions. However, the starch seen for K2C was minimal (Supplemental Figure 1). Considering that K6C represents an even more deprived nitrogen to carbon ratio compared to K2C, it was anticipated that K6C would show the lowest levels of starch since both conditions also show significant lipid accumulation. However, the increase in starch content for K6C suggests that these cells still have abundant carbon available for transformation into carbohydrates, and potential future engineering efforts could focus on knocking out genes associated with starch accumulation to divert this excess energy toward lipid feedstock production instead.

The confocal imaging also happened to capture a possible lipid droplet secretion event. During live-cell imaging, a single cell was captured over a few minutes showing the formation of a cellular bleb containing a single lipid droplet that was released in subsequent scans (Supplemental Figure 2). We have previously reported that this organism does not appear to have any canonical proteins associated with lipid droplet secretion from other organisms^22^. That study also captured static images showing what was described as blebbing intermediates. The current image series reported here is the first case in which the process was observed live thereby lending additional support to the theory that *O. tauri* is capable of lipid droplet secretion although the detailed mechanism remains elusive.

Dramatic changes observed for lipid staining profiles prompted the exploration of underlying global proteomic expression profiles for each experimental culture condition. Cultures were harvested at 24- and 48-hour time points for LC-MS/MS proteomics to obtain a measure of global proteomic expression. These time-points were chosen for comparison since they showed the biggest relative 24-hour changes via FACS and confocal analysis. The proteomics data was mapped into individual metabolic groups (synthesis, degradation, energy, other and non-metabolic pathways) to interpret relative changes of cells cultured in K6CN, K2CN, K2C and K6C conditions for each metabolic pathway (Fig. 2). Of particular note was the increased changes in abundance for both K2C and K6C in the carbohydrate synthesis pathway compared to K2CN and K6CN. However, very few proteins within the fatty acid (FA)/lipid synthesis pathways showed significant, if any, change in abundance despite bioimaging observations of cellular lipid increases up to 60% the volume of cells. For example, while DGAT, the canonical enzyme representing the last committed step of TAG synthesis was detected with global proteomics, it’s abundance remained effectively equivalent across all 4 sample conditions.

**Figure 2.**
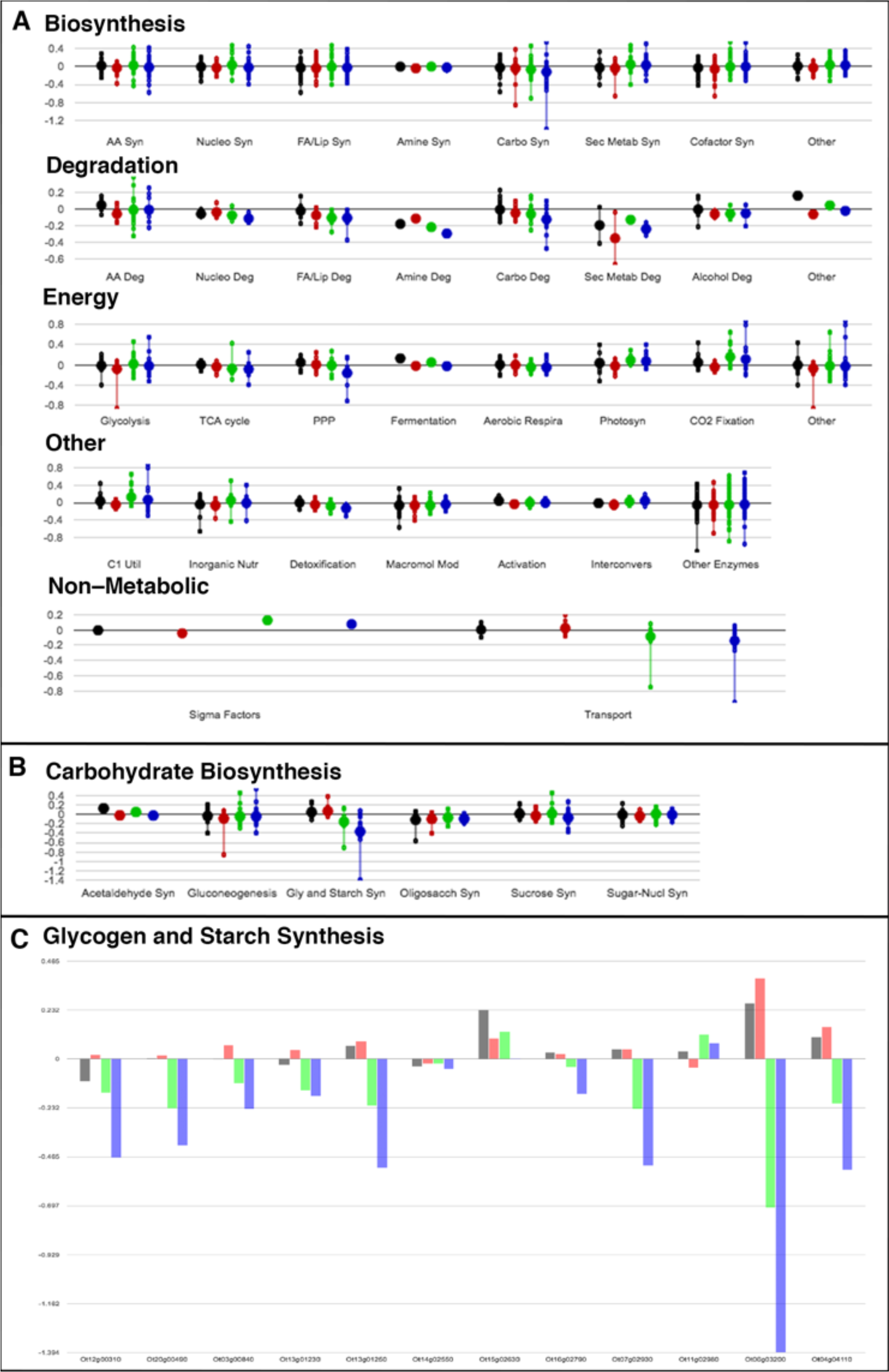
Metabolic pathway mapping of global proteomics. *O. tauri* cells cultured in K6CN (black), K2CN (red), K2C (green), K6C (blue) at 48 hours relative to cultures from K2CN at 24 hours were grouped and mapped by metabolic pathway(A), subpathway (B), and individual proteins (C). Each large circle represents an average level of abundance on a log2 scale where negative values are increases in abundance relative to cell conditions K2CN at 24 hours. Smaller circles represent individual proteins in the respective metabolic pathway.

High C to N ratio of K6C displayed consistently higher proteomic responses compared to K2C for upregulated proteins related to C storage. The complete starch pathway was detected with ascending upregulation from alpha amylase to granule-bound starch synthetase (GBSSI) for N depletion conditions. GBSSI was the third highest upregulated protein overall highlighting the reliance on carbon storage under these conditions. Previous studies have found similar results for GBSSI^23^. N deprivation also caused downregulation of proteins involved in N acquisition, such as nitrate transporters, nitrate reductase with concomitant upregulation of N scavenging proteins for glutamine, asparagine, and urea. Much of the downregulated proteins detected were ribosome based or were proteins localized to the chloroplast.

Overall 471 unnamed protein products were found to be upregulated or downregulated with several exhibiting some of the most extreme abundance changes (Fig. 4). N deplete conditions K2C and K6C had similar distributions of upregulated and downregulated protein trends, with more increased upregulation of proteins for excess carbon cultures. For K6C, 5 of the 10 most upregulated and 3 of the 10 most downregulated were UPP (Fig. 3). Interestingly, several of these proteins show inversion of abundance for replete versus depleted N conditions. XP_003075209 (ostta02g03680), XP_003081059 (ostta09g00670), XP_003078347 (ostta03g04500) all exhibit significant upregulation under K6CN and K2CN conditions at 48 hours compared to K2CN at 24 hours, however, these proteins show significant downregulation for K2C and K6C conditions at 48 hours. The biggest change was seen for XP_003078347 (ostta03g04500) which was had a log2 value change of −0.54 for K6CN but a value of 1.96 for K6C. Similarly, other unnamed proteins XP_003084215 (ostta18g01710), XP_003082140 (ostta11g03180) and XP_003082699 (ostta13g02170) were all downregulated for K6CN and K2CN at 48 hours but upregulated for K2C and K6C. Clearly these unnamed proteins are dramatically affected by the bioavailable ratio of C and N and represent interesting targets for future in-depth functional annotation.

**Figure 3.**
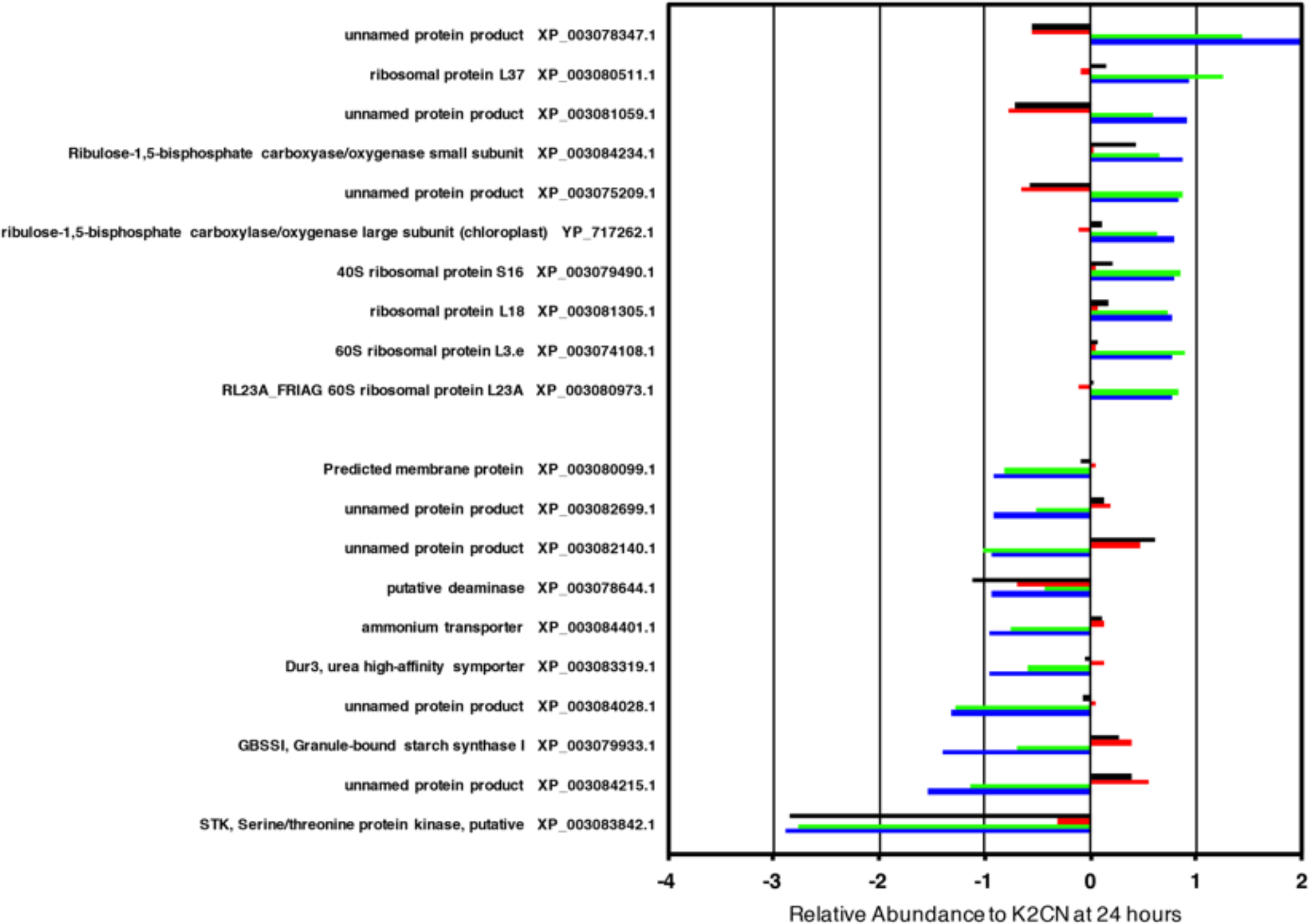
Selection differentially expressed proteins from global proteomics of varying N and C conditions. Abundance profiles for 10 most downregulated (positive values) and upregulated (negative values) proteins comparing K2CN at 24 hours versus K6CN at 48 hours (black), K2CN at 48 hours (red), K2C at 48 hours (green), and K6C at 48 hours (blue) cell cultures.

**Figure 4.**
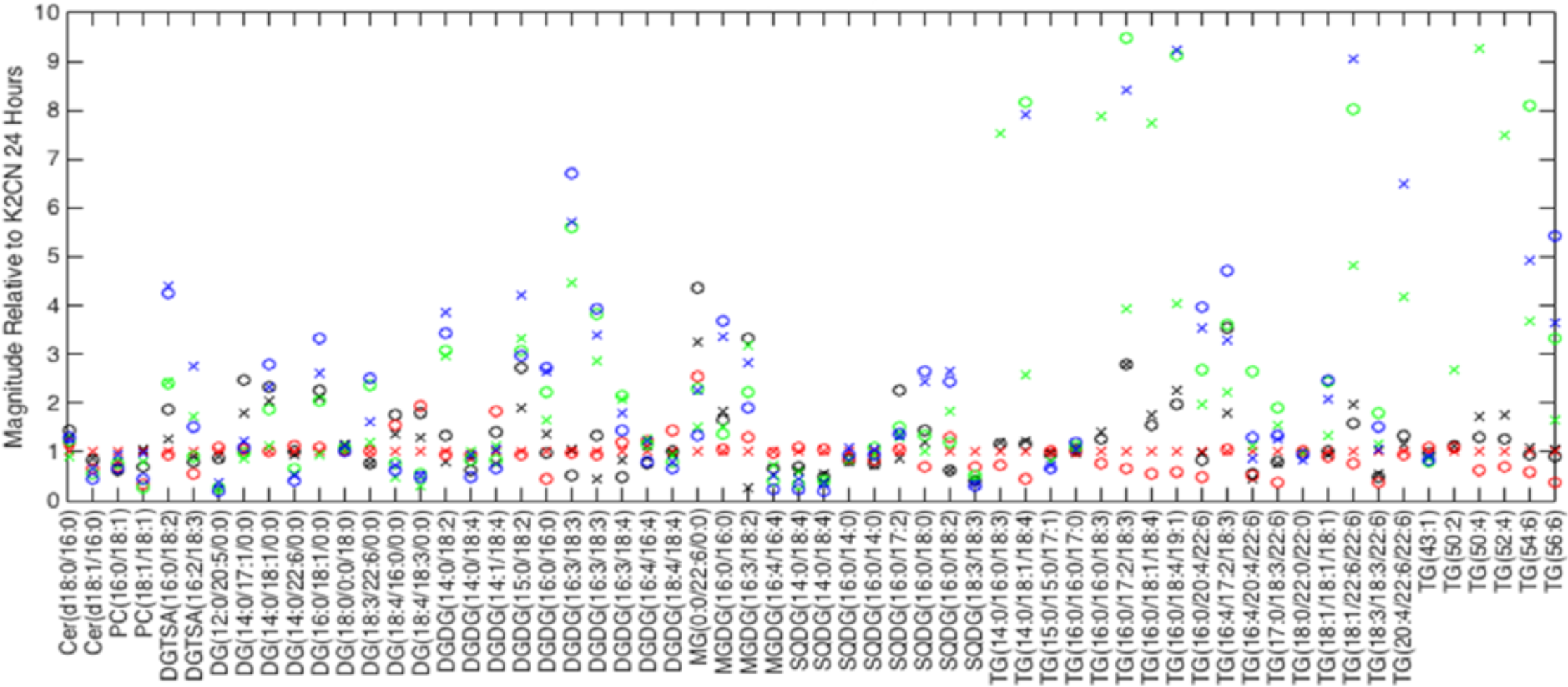
Time resolved differential expression of global lipids. Selection of lipids (out of 287) detected via LC-MS/MS for cell cultures at 24 (X symbols) and 48 hours (O symbols) for varying C:N ratio conditions K6CN (black), K2CN (red), K2C (green) and K6C (blue). *Cer: Ceramide; PC: phosphatidylcholine; MGDG: monogalactosyldiacylglycerol; DG: diacylglycerol; DGDG: digalactosyldiacylglycerol; DGTSA: diacylglyceryl trimethylhomoserine or diacylglyceryl trimethyl-beta-alanine; SQDG: sulfoquinovosyldiacylglycerol; and TG: triacylglycerols*

Conducting a BLAST alignment analysis on the UPP identified in these runs resolved a predicted membrane protein (XP_003080099 (ostta06g04530)) upregulated for N depleted conditions K2C and K6C, which was identified as having a domain with homology to TMEM14, an uncharacterized superfamily believed to be involved in membrane transport of lipids in higher eukaryotes. This protein could possibly play a role in the lipid droplet secretion from *O. tauri*. Other potentially interesting unknown protein products were also uncovered during BLAST alignments with two notable probable identifications: A putative Acyl-CoA N-acyltransferase (XP_003084085.1 (ostta18g00460)) slightly upregulated for K2CN and K2C but not K6CN or K6C; and a putative Zinc finger (XP_003080741.1 (ostta08g01730)) slightly downregulated for K6CN, K6C, and K2C conditions, which have been known to participate in a number of eukaryotic cellular mechanisms including lipid binding. Additional proteins such as sarcosine-dimethyltransferase (SDMT), an enzyme found in the betaine biosynthesis pathway in other algae and higher plants related to osmoprotection during cellular stress^24,25^, were found to be elevated only for K2C and K6C conditions, which may reinforce cellular stability during N stress and lipid accumulation.

Despite observing limited changes in the proteome related to lipid metabolism, the visual confocal and FACS analysis provided ample evidence of significant lipid accumulation that warranted further characterization to understand cellular lipid composition and relative abundance. Thus, the same K6CN, K2CN, K2C, and K6C cultures were surveyed at the same 24- and 48-hour time-points using LC-MS/MS global lipidomics analysis. Quantitative lipid profiles were collected for more than 280 lipids (Supplemental Figures 3 – 9). The N depletion conditions (K2C and K6C) resulted in significant increases of energy dense TAG lipids with more than 10 TAGs showing >40-fold increase in abundance already at 48-hours. The FACS and confocal analysis showed that the NL content continues to accumulate well beyond 48 hours. In comparison, TAG lipid abundance was flat or decreased for K6CN and K2CN conditioned cells, indicating that cells were not diverting C into lipid energy storage during nutrient rich conditions although they were still accumulating starch as seen from SRS (Supplemental Figure 1). The observed FA profiles provide additional support for recently reported^26^ unique long chain FAs despite lacking annotated enzymes known to synthesize them, indicating that *O. tauri* is at the very least an intriguing oleaginous organism for lipid feedstock development. In addition, the number of long chain FAs in TAGs were significantly enhanced during K6C conditions, demonstrating this organism’s capacity to uptake and transform excess C into long chain energy dense lipids without genetic modification (Supplemental Figures 7–9).

Changes in structural lipids were also detected (Supplemental Figures 3 – 6). Thylakoidal membranes are composed of monogalactosyldiacylglycerol (MGDG) and digalactosyldiacylglycerol (DGDG), which stabilize the thylakoid for maximal photosynthetic efficiency ^27^. In addition, algal thylakoid membranes contain abundant amounts of sulphoquinovosyldiacylglycerol (SQDG) which contributes to increased stability in the photosynthetic harvesting complexes and accommodates membrane protein associations unique to microalgae ^28^. Increased abundances for MGDG and 3- to 7-fold increases in DGDG were detected for K2C and K6C conditions, with C16 chain lengths exhibiting the most abundant and dramatic changes during N depletion. C16 chain length SQDG lipids were also slightly upregulated for K6C, suggesting they may be adding stability to light harvesting complexes or membrane protein expression unique to excess C exposure and N depletion.

## Discussion

Linkages between N and C metabolism related to lipid biogenesis were interrogated for the microalgae *Ostreococcus tauri*. Single cell and population imaging experiments combined with global proteomic and lipidomic experiments on the same cultures demonstrated that even the simplified cell architecture and genome of *O. tauri* displays complicated regulatory linkages as a function of bioavailable carbon to nitrogen ratios. Cryogenic soft x-ray nanotomography revealed distinct lipid droplet distributions of varying sizes confirming initial fluorescence microscopy results. K2C conditioned cells had uniform sized lipid droplets whereas cells in K6C conditions had lipid droplets of varying size including very large lipid droplets that swelled and deformed the cells.

Numerous photosynthetic, structural, and energy storage fatty acid (FA)/lipids were verified through LC–MS/MS lipidomic analysis that included detection of long chain TAGs, ideal for lipid feedstocks. Proteomic data combined with physiological cell responses to varying C and N revealed that although FA/lipid synthesis pathways had little proteomic changes, carbohydrate synthesis proteomics was independently upregulated. Both ribulose 1,5-biphosphate carboxylase/oxygenase large and small subunits (Rubisco) were slightly downregulated for K2C and K6C relative to K2CN suggesting Rubisco expression in *Ostreococcus* contains minimum concentrations of Rubisco to support normal growth as reported for other organisms^29^. Restricting protein identifications to subsets related to carbohydrate conversion, N scavenging, and energy regulation allowed for the simplification of biological interpretations related to lipid feedstock optimization targets. The most upregulated protein was a protein kinase involved in serine/threonine phosphorylation and was equally upregulated for K2C, K6C, and K6CN conditions, which could possibly play a role in regulating diurnal cycling or stress response since K6CN could also be seen as another N deprived state relative to available C. GBSSI was the third most upregulated protein in all and was more upregulated in K6C than K2C. Knock-down of GBSSI could potentially provide a redirection of *O. tauri* metabolism to less starch accumulation and more lipid accumulation under K6C conditions.

In addition to the proteins related to energy storage, increased abundances of proteins related to lipid viability, lipid production, and osmotic shock were detected to varying degrees. Finally, a surprising number of proteins of unknown function and identity were revealed to be part of the 10 most significant proteomic increases and decreases overall and each is a potential target for future engineering efforts. A recent study focused on circadian protein regulation^30^ identified ostta03g04500, ostta09g00670, and ostta02g03680 proteins of unknown function that happen to overlap with the findings reported here. This suggests that the proteins found in both studies may have a co-relational effect between carbon and nitrogen bioavailability and the circadian cycle. Delving further into the function of these individual proteins will be important for understanding their real role in overall cellular regulation and metabolic processes. Furthermore, the wealth of undefined proteins emphasizes the extent of unexplored opportunities related to N, C, and energy storage pathways, and highlights the peculiar genetic diversity within natural populations of *Ostreococcus* and possibly other primitive marine species.

## Methods

### Strain maintenance, culture growth media, and nutrient starvation conditions

*O. tauri* cell cultures were obtained from the Roscoff Culture Collection (RCC745); strain name: OTTH0595, which has been fully sequenced ^20^. Cultures of RCC745 were grown in defined Keller (K) media^31^ with normal or depleted N and HCO_3–_: K2CN contained normal N and C, K6CN with normal N and 6mM HCO_3–_, K2C with depleted N and 2.5mM HCO_3–_, and K6C with depleted N and 6mM HCO_3–_. All Keller media-based culture conditions were prepared in fresh artificial seawater (ASW) with defined amounts of nutrients analytically prepared fresh and sterile filtered prior to each experiment. To monitor growth and lipid accumulation over time absorbance at 680nm and 750nm was measured to obtain both values for chlorophyll content and particulate matter, respectively, for each culture. Graphing the ratio of 750nm/680nm provided a measure of chlorophyll functional efficiency as well as possible lipid particulates in solution. Graphical analysis of each growth and starvation curve required minimal normalization due to our consistent efforts in capturing cells during mid-log stages of growth. For growth and lipid accumulation studies cells were grown initially in normal K media to approximately 0.03 at OD680 then gently centrifuging cultures at 1200XG for 10 minutes in a swing bucket rotor, washed with respective defined K media, and resuspending cells into defined K media at a target 0.03 at OD680 and continued diurnal light entrainment for specified time courses in sealable CytoOne non-treated cell culture flasks (USA Scientific, USA) with mixing of cultures once per 24 hours. To prepare cell cultures for starvation surveys they were gently centrifuged fresh cultures at 2200xG for 10 mins with swing bucket rotor centrifugation and washed cell pellets once with defined K media of interest then suspended cells in defined media conditions and continued diurnal light entrainment for specified time courses in sealable CytoOne non-treated cell culture flasks (USA Scientific, USA) with mixing of cultures once per 24 hours.

### Fluorescence activated cell sorting analysis of intracellular lipid content

*O. tauri* cells were cultured to mid log phase and gently centrifuged to concentration then stained with Nile Red (4.8*μ*g/mL) lipid stain for exactly 10 mins before each experimental measurement on the BD INFLUX flow cytometer (BD Biosciences, San Jose, CA, USA). FSC and SSC were used to gate out any non-specific cellular debris. Specific gating in the range of known cell size of *O. tauri* was used to determine the fluorescence from stained neutral lipid (488/542±13.5 nm), phospholipid (561/615±12 nm), and natural chlorophyll autofluorescence (640/670±15 nm) for defined populations of cells. Each individual FACS experiment was calibrated to 3.6 ide scatter 10 mins before running our sample measurements in defined media cultures. The fluorescence intensity of neutral lipid and phospholipid fluorescence intensity at specific time points was compared in scatter plots to demonstrate population dynamics for each sample condition. K2CN at 24 hours was used as the baseline for normal conditions to detect changes due to varying C:N bioavailability.

### Fluorescence and SRS confocal microscopy

Confocal images were obtained on Zeiss LSM 710 (Carl Zeiss AG, Germany) confocal microscope with a 100x oil immersion objective. An InTune Laser with 505nm and 535nm light was used to maximize the separation of the triglyceride (585nm) and phospholipid (638nm) emission peaks while diminishing crosstalk of the Nile Red stained cells. In addition, chlorophyll autofluorescence was excited with 405nm light and monitored the emission profile at 680nm. Nile Red stained cells were immobilized on glass slides with poly-L-lysine and imaged immediately with z-scan slicing of 0.43*μ*m to survey whole cell fluorescence labeling distribution. All fluorescence channels were set with identical gain and laser power settings to provide relative levels of fluorescent intensity and no adjustments of contrast or gain were applied to fluorescence imaging during post processing. *O. tauri* cultures were grown in 15mL falcon tubes in a 12:12 light:dark illumination (~20 *μ*mol/m^2^/s) at 25ºC for 96 hours, harvested by gentle centrifugation (1000xG, 5 mins) and concentrated equally to 10x original cell density for imaging. Concentrated cell suspensions were then stained with Nile Red (4.8 *μ*g/mL) for 10 minutes and then 3*μ*L of the labeled culture were mounted on poly-lysine treated microscope slides (Electron Microscopy Sciences, USA) for confocal fluorescence and SRS imaging. Fluorescence and SRS confocal microscopy were conducted on the Nile Red stained *O. tauri* cells using a Leica DMi8 (Leica Microsystems GmbH, Germany) inverted confocal microscope (a 63x/1.20na water immersion objective) integrated with an APE picoEmerald laser consisting of a 2.5 ps pulsed tunable pump and 1031nm stokes with the SRS detection module and an EOM Modulator. Prior to the SRS detection, the stokes beam was blocked by a FESH1000 filter (Thorlabs, USA). The SRS signal was detected on a 10x10mm photodiode connected to the APE lock-in amplifier. For starch detection the SRS pump signal was tuned to 947nm at 152mW with bandwidth of 0.7nm with the stokes power at 150mW modulated by the integrated EOM, and a delay of 3500fs to detect the 860cm^‒1^ peak of starch ^32^. Cellular chlorophyll was excited with a 638nm laser with 1% power and fluorescence signal collected with emission range of 677–689nm. Neutral lipid fluorescence from Nile Red staining was excited with 552nm with 1% laser power and emission 575–583nm. Sequential acquisition of different channels was acquired at 1024x1024 format, with slower 400 scan speeds, and 4x zoom. Maximum thresholds were changed consistently for all SRS starch images to obtain equal post processing values across all samples; no post processing was conducted on either chlorophyll or neutral lipid channels.

### Cryogenic Soft X-ray Tomography for Intracellular Distribution of Organelles

Cell cultures were grown on-site at the Advanced Light Source (ALS) at Lawrence Berkeley National Laboratory with a homemade 470nm light source with a measured intensity of 20**μ** E. Cell cultures were incubated with 12-hour diurnal light at RT and harvested by centrifugation at 48, 72 and 96-hour time points. Cells were gently centrifuged at 1000xG for 10mins to pellet. Pelleted cells had all but ~5**μ** L of supernatant removed to remove a viscous cell biomass. Wet cell biomass was inserted into 5**μ** m micro capillaries and plunge frozen in cold liquid propane. In some cases, 6**μ** m polystyrene beads were added to cell suspensions prior to centrifugation to minimize the impact of freezing on large lipid containing cells. Frozen capillaries were stored in liquid nitrogen until imaging by the soft x-ray light source. Soft x-ray data acquisition was carried out on beamline 2.1, a soft x-ray microscope in the National Center for X-Ray Tomography (NCXT) located at the Advanced Light Source in Berkeley, California^33^. The microscope soft x-ray illumination was generated by a bend-magnet in the synchrotron lattice and focused onto the specimen by a Fresnel Zone Plate (FZP) condenser. Specimen illumination was order-sorted by a pinhole positioned just in front of the specimen. A second zone plate, located downstream of the specimen, magnified and focused an image of the specimen on a CCD detector. During data collection, the cells were maintained in a stream of helium gas that had been cooled to liquid nitrogen temperatures. Each tomographic dataset (i.e., 90 projection images spanning a range of 180°) was collected using Fresnel zone plate based objective lens with a resolution of 50 nm. Exposure times for each projection image ranged from 150 to 300 msec. The software suite AREC3D was used to align the projection images calculate tomographic reconstructions^34^.

### Lipid Extraction and Lipidomics Using Liquid Chromatography Tandem Mass Spectrometry

Each *O. tauri* culture was harvested at specific time points and ultimately spun down at 3000 x g for 10 min to form a pellet in chloroform compatible 2 mL Safe Seal microfuge tubes (Sorenson Bioscience, Inc, Salt Lake City, UT). The supernatant was removed, and the wet pellet was weighed for an estimate of biomass, then a mixture of 1 mL of methanol, 0.5 mL of chloroform and 0.4 mL of nanopure water was added to each pellet. The pellets were vortexed and sonicated for about 10 sec in a bath sonicator and the mixture was allowed to stand at room temperature for 10 min. Then an additional 0.5 mL of chloroform and 0.5 mL of water were added, and the mixtures were shaken vigorously into an emulsion. Each sample was centrifuged at 10,000 x g for 10 min. The bottom chloroform layer was carefully removed from the separated solvent layers using a Pasteur pipette and taking care not to take up any of the protein disc or the top polar layer and was placed in a pre-weighed glass auto sampler vial. The bottom lipid layer was dried down overnight under nitrogen and the vials with the total lipid extract (TLEs) were weighed to obtain the total mass. 5*μ*L of chloroform was added to each dried sample and then the sample diluted to 50 *μ* g/*μ*L with methanol. The samples were then stored at –20 °C until ready for mass spectrometric analysis.

Extracted lipids were dried down then reconstituted in methanol. The TLEs were analyzed by LC-MS/MS using a Waters NanoAquity UPLC system interfaced with a Velos Orbitrap mass spectrometer (Thermo Scientific, San Jose, CA) as outlined in Dautel et al. 2017^35^. LC-MS/MS raw data files were imported into the in-house developed software LIQUID^36^ for identification of lipid molecular species. Manual validation of the lipid identifications were determined by examining the tandem mass spectra for diagnostic ion fragments along with associated chain fragment information. In addition, the isotopic profile, extracted ion chromatogram (XIC), and mass error of measured precursor ion were examined for lipid

### Cellular Protein Extraction and Digestion

To the remaining protein/debris pellet obtained from the lipid extraction, 1 mL of ice cold methanol was added 3 x to wash the majority of residual metabolites away from the pellet. A Filter-Aided Sample Prep (FASP) protein extraction and trypsin digestion was then performed using the FASP protein digestion kit (Expedeon, San Diego CA) using manufacturer’s suggested protocol (Erde J et al, 2014, J Proteome Res 13(4):1885-1895). Briefly, methanol washed pellets were air dried for 2 hrs and were resuspended in solubilization buffer of 12 mM deoxycholate, 12 mM N-lauroyl sarcosine with 10 mM TCEP, and 200 mM ammonium bicarbonate, pH 8.0 at an approximate concentration of 13 *μ*g/*μ*L. 30 *μ*L of this protein solution was then mixed with 200 *μ*L of urea sample solution (kit provided), the sample was centrifuged on the kit provided spin filter at 14,000 x g for 15 min. Washes with urea and ammonium bicarbonate along with trypsin digestion and alkylation with iodoacetamide were carried out as the kit specifies. Peptides were then suspended in nanopure water and peptides were then quantified using a BCA assay (Pierce, Rockford IL) with a bovine serum albumin standard.

### iTRAQ Peptide Labeling

Peptides were labeled with 8-plex iTRAQ (AB Sciex, Redwood City, CA) reagents as described below. 100 *μ*g of each peptide sample was placed in a new tube and dried down. Chanel designations are as follows: 43 *μ*g of dissolution buffer (iTRAQ buffer kit) was added to each sample, these being vortexed into solution and centrifuged briefly to draw sample to the bottom of each tube. The iTRAQ reagent (30 *μ*L) was diluted further with isopropanol (115 *μ*L) and this was then added to each sample. Each reaction was carried out at RT for 2 hrs, with 50 mM ammonium bicarbonate (200 *μ*L) added to quench each reaction tube. After 1 hr, the contents from all iTRAQ channel reactions were added to one tube and then the sample was vortexed and dried down in a speed vac. The labeled peptides were cleaned up using C-18 SPE columns (SUPELCO Discovery) were then employed to remove the salts, using a 0.1% TFA in nanopure water to wash the peptides and 80% acetonitrile, 0.1% TFA in water to elute the peptides.

#### Offline Fractionation of Peptides and Preparation of Proteome Samples

400 *μ*g of iTRAQ labeled peptides were separated using an off-line high pH (pH 10) reversed-phase (RP) separation with a Waters XBridge C18 column (250 mm x 4.6 mm column containing 5 *μ*m particles and a 4.6 mm x 20 mm guard column) using an Agilent 1200 HPLC System. The sample loaded onto the C18 column was washed for 15 min with Solvent A (10 mM ammonium formate, adjusted to pH 10 with ammonium hydroxide). The LC gradient started with a linear increase of Solvent B (10 mM ammonium formate, pH 10, 90% acetonitrile in water) to: 5% over 10 min, 45% Solvent B over 65 min, and then a linear increase to 100% Solvent B over 15 min. Solvent B was held at 100% for 10 min, and then was changed to 100% Solvent A, this being held for 20 min to recondition the column. The flow rate was 0.5 mL/min. A total of 96 fractions were collected into a 96 well plate throughout the LC gradient. The high pH RP fractions were then combined into 12 fractions using the concatenation strategy previously reported^37^. Peptide fractions were dried down and re-suspended in nanopure water at a concentration of 0.075 *μ*g/ *μ*L for mass spectrometry analysis using a Q Exactive HF Hybrid Quadrupole-Orbitrap MS (Thermo Scientific) system as described below.

### Mass-Spectrometry Based Analysis of Samples

All peptide samples were analyzed using an automated home-built constant flow nano LC system (Agilent) coupled to an Q Exactive HF Hybrid Quadrupole-Orbitrap MS (Thermo Fisher Scientific). Electrospray emitters were custom made using 150 *μ*m o.d. × 20 *μ*m o.d. × 20 *μ*m i.d. chemically etched fused silica. An on-line 4-cm × 360 *μ*m o.d. × 150 *μ*m i.d. fused-silica capillary analytical column (3 *μ*m Jupiter C18) was used. Mobile phases consisted of 0.1% formic acid in water (A) and 0.1% formic acid acetonitrile (B) operated at 300 nL/min with a gradient profile as follows (min:%B); 0:5, 2:8, 20:12, 75:35, 97:60, 100:85.

### Peptide Identification, Quantification and Analysis

For peptide identification, MS/MS spectra were searched against a decoy O. tauri database using the algorithm SEQUEST. An approach to correlate tandem mass spectral data of peptides with amino acid sequences in a protein database^38^. Search parameters included: no enzyme specificity for proteome data and trypsin enzyme specificity with a maximum of two missed cleaves, ± 50 ppm precursor mass tolerance, ± 0.05 Da product mass tolerance, and carbamidomethylation of cysteines and iTRAQ labeling of lysines and peptide N-termini as fixed modifications. Allowed variable modifications were oxidation of methionine. MSGF+ spectra probability values were also calculated for peptides identified from SEQUEST searches^39^. Measured mass accuracy and MSGF spectra probability were used to filter identified peptides to <0.4% false discovery rate (FDR) at spectrum level and <1% FDR at the peptide level using the decoy approach. iTRAQ reporter ions were extracted using the MASIC software^40^ for fast quantitation and flexible visualization of chromatographic profiles from detected LC-MS(/MS) features with a 10-ppm mass tolerance for each expected iTRAQ reporter ion as determined from each MS/MS spectrum.

Relative abundances of peptides were determined using iTRAQ reporter ion intensity ratios from each MS/MS spectrum. Individual peptide intensity values were determined by dividing the base peak intensity by the relative ratio associated with each reporter ion. All peptide values were then transformed into log2 values for comparison between conditions.

## Acknowledgements

This research was performed using the Environmental Molecular Sciences Laboratory (EMSL), a national scientific user facility sponsored by the Department of Energy’s Office of Biological and Environmental Research and located at PNNL. This research also used resources of the Advanced Light Source, which is a DOE Office of Science User Facility under contract no. DEAC02-05CH11231.

## Contributions

JEE devised and managed all experiments with input from CRS. CRS conducted confocal and Raman imaging experiments and coordinated integrated omics analysis. KKH and CDN performed lipid and proteomics sample prep. RM collected proteomics data that SP analyzed, and NK and WRC worked into the cell model. JEK performed lipidomic data collection and lipidomic analysis. Cryogenic soft X-ray sample preparation was conducted by JHC, RB and CRS with JHC and GM acquiring tilt series for subsequent reconstruction by AE. FACS analysis was conducted by WC. JEE and CRS wrote initial manuscript and all authors edited and approved final text.

---

**Figure S1.**
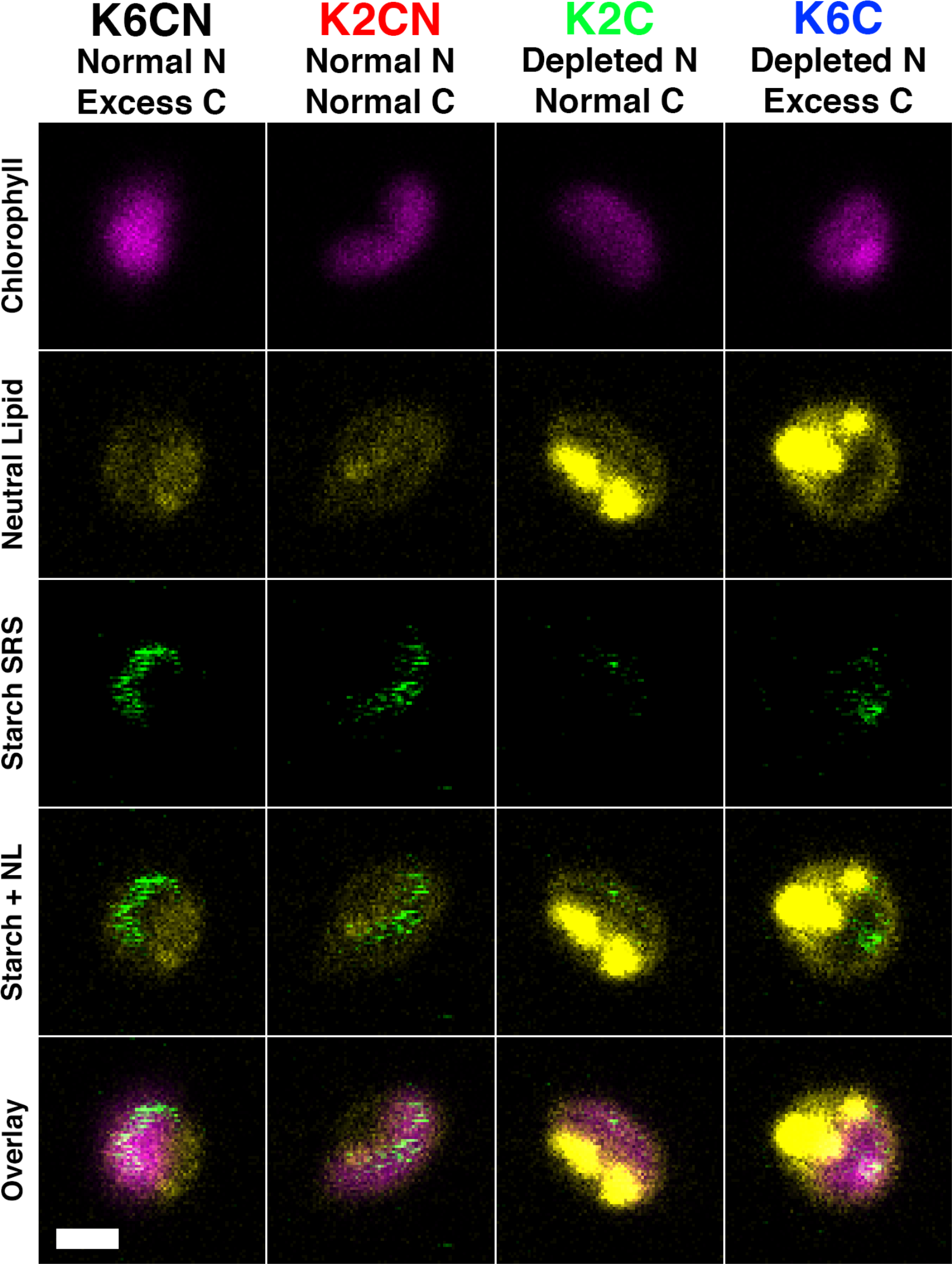
Fluorescence and SRS verification of neutral lipid and starch accumulation. Fluorescence and SRS microscopy were conducted on *O. tauri* cells stained with Nile Red from various C:N conditions after 96 hours of starvation. Chlorophyll autofluorescence (pink) was observed in all cells with neutral lipid (yellow) and SRS starch signal (green). Elevated neutral lipid was observed in K2C and K6C cells localized outside the chloroplast. Starch was detected and localized to the chloroplast for K6CN, K2CN, and to a lesser extent K6C, with little to no starch signal found in K2C cells. Scale bar represents 1*μ*m.

**Figure S2.**
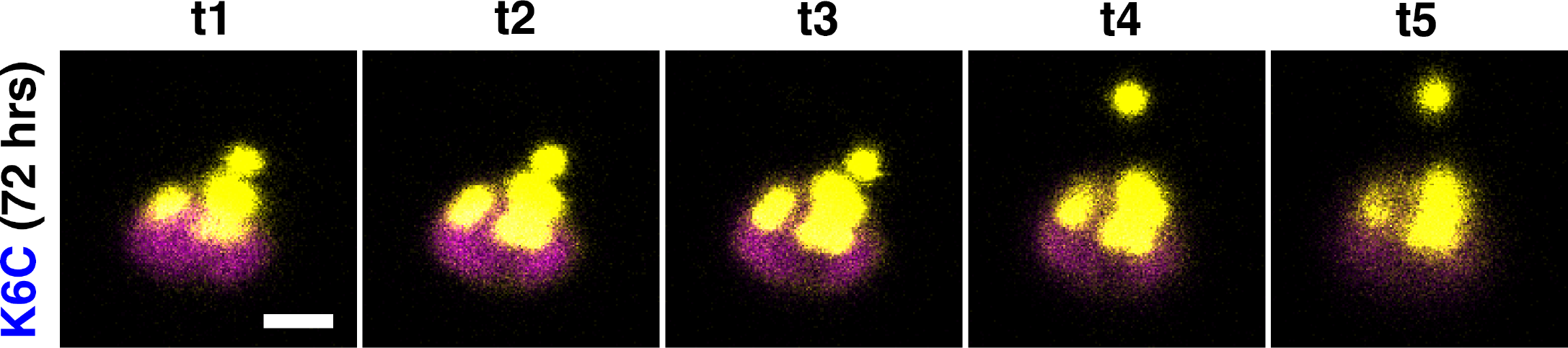
Time course of lipid release from cells into intercellular space. *O. tauri* cells cultured in K6C media for 72 hours were harvested and stained with Nile Red and then imaged by confocal fluorescence microscopy. Chlorophyll autofluorescence (pink) and neutral lipid (yellow) revealed many large lipid bodies during z-slice imaging and a single lipid body release was tracked over time (t) in intercellular milieu while other lipids remained inside the intracellular space.

**Figure S3.**
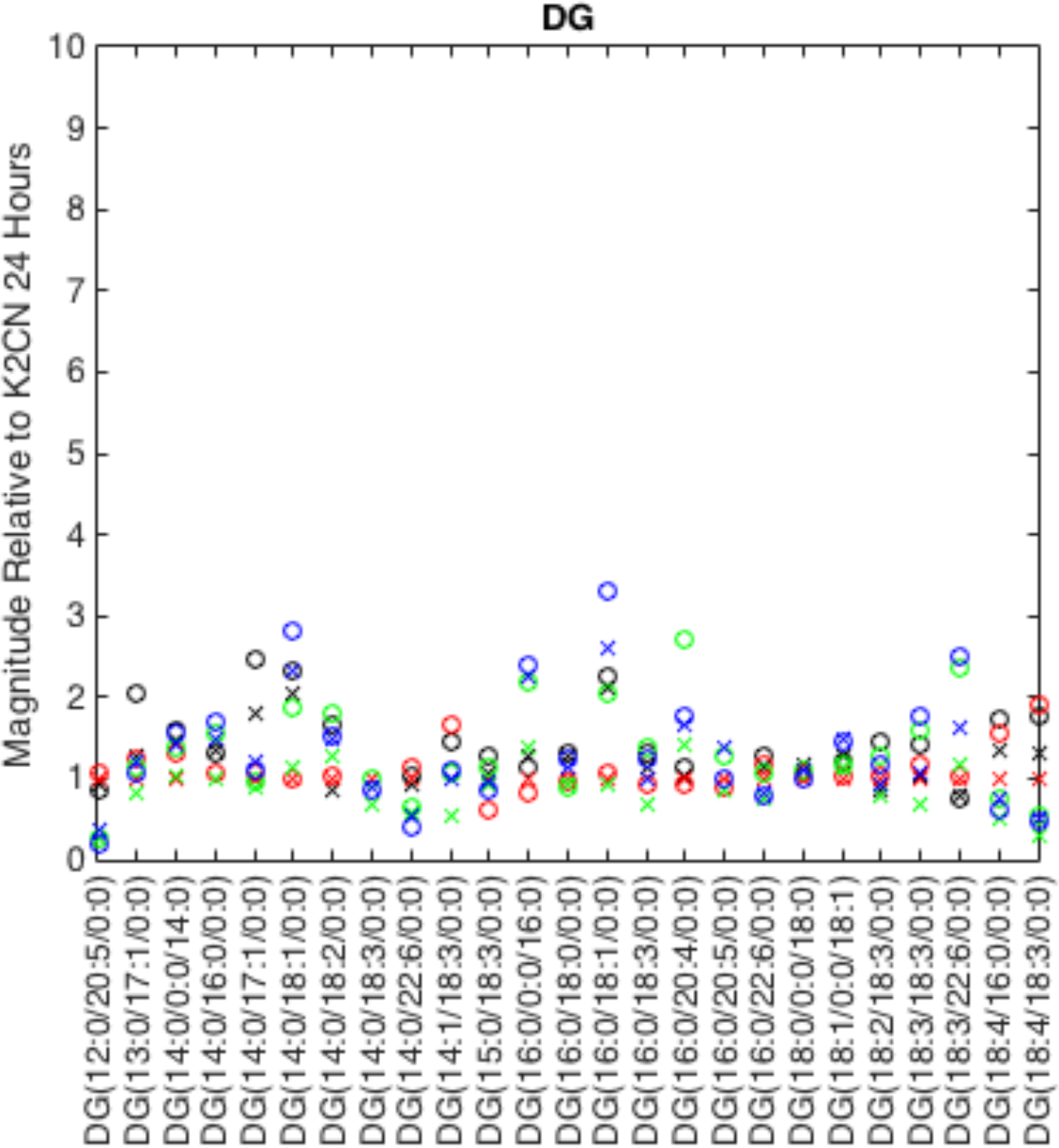
Time-resolved lipidomics of diacylglycerol lipids. *Diacylglycerol* (DG) lipids detected via LC-MS/MS for cell cultures at 24 (X symbols) and 48 hours (O symbols) for varying C:N ratio conditions K6CN (black), K2CN (red), K2C (green) and K6C (blue) relative to K2CN at 24 hours (y-axis).

**Figure S4.**
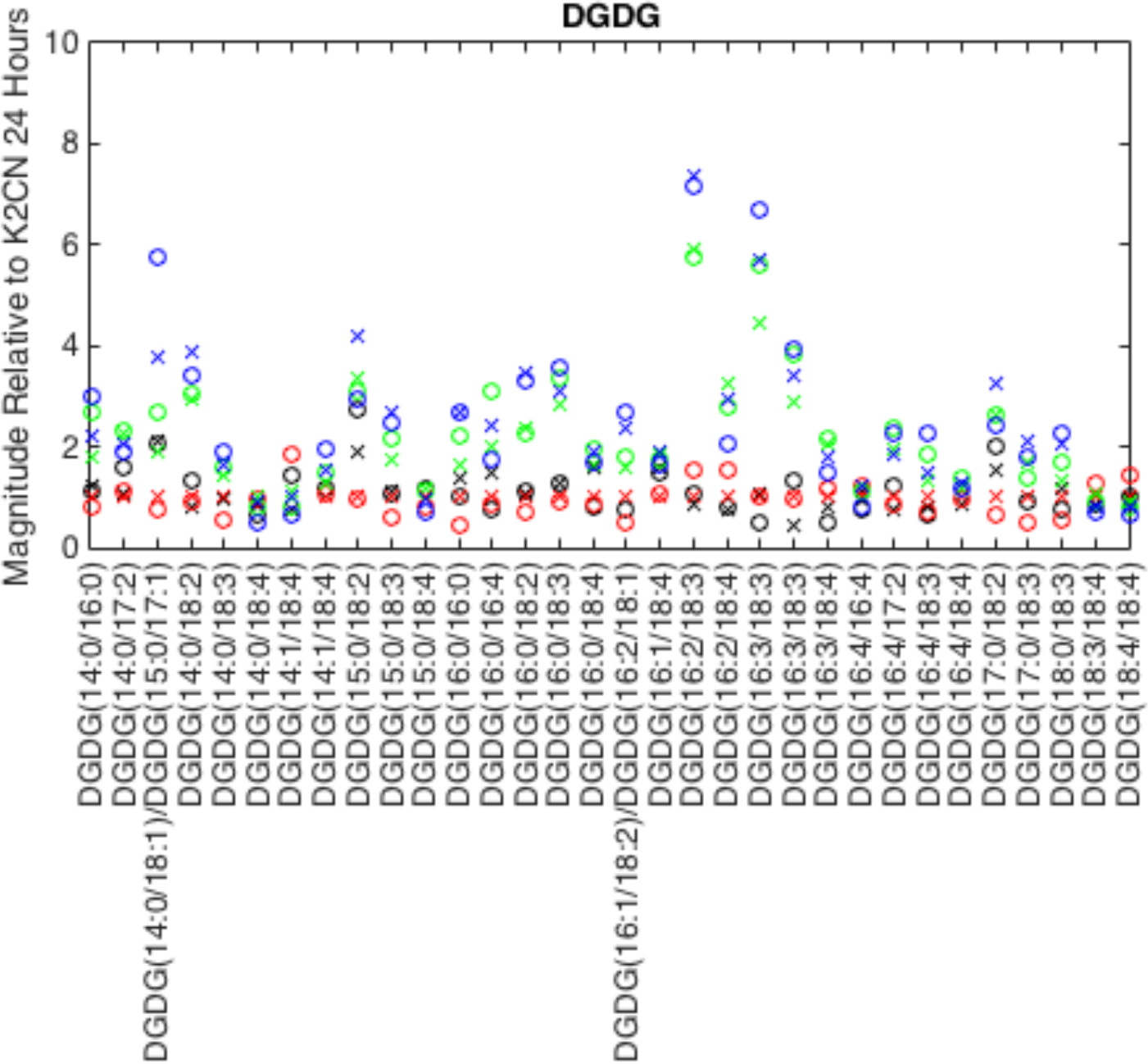
Time resolved differential expression of digalactosyldiacylglycerol lipids. Digalactosyldiacylglycerol (DGDG) lipids detected for cell cultures at 24 (X symbols) and 48 hours (O symbols) for varying C:N ratio conditions K6CN (black), K2CN (red), K2C (green) and K6C (blue) relative to K2CN at 24 hours (y-axis).

**Figure S5.**
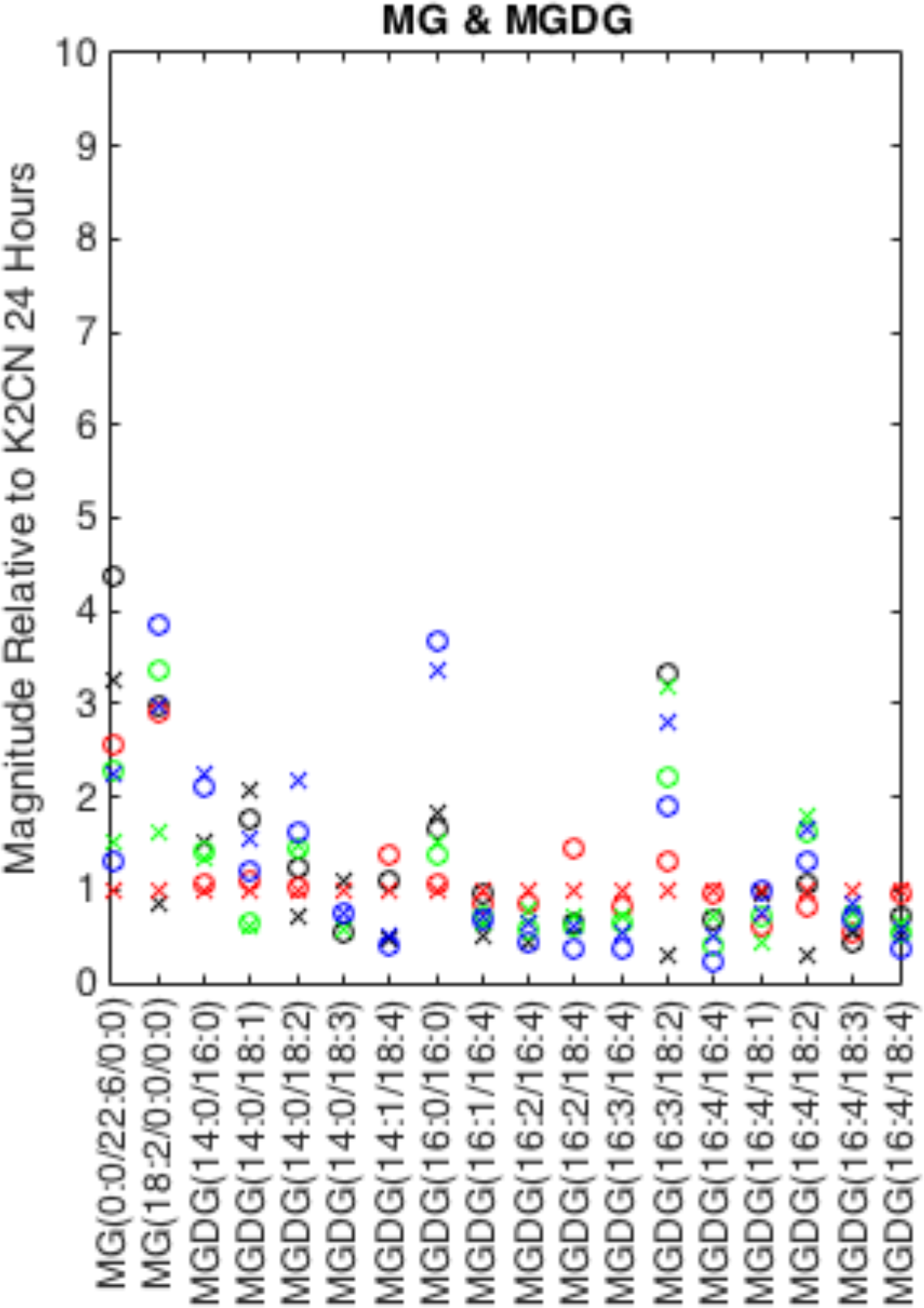
Time resolved differential expression of monoacylglycerol and monogalactosyldiacylglycerol lipids. Monoacylglycerol (MG) and monogalactosyldiacylglycerol (MGDG) lipids detected via LC-MS/MS for cell cultures at 24 (X symbols) and 48 hours (O symbols) for varying C:N ratio conditions K6CN (black), K2CN (red), K2C (green) and K6C (blue) relative to K2CN at 24 hours (y-axis).

**Figure S6.**
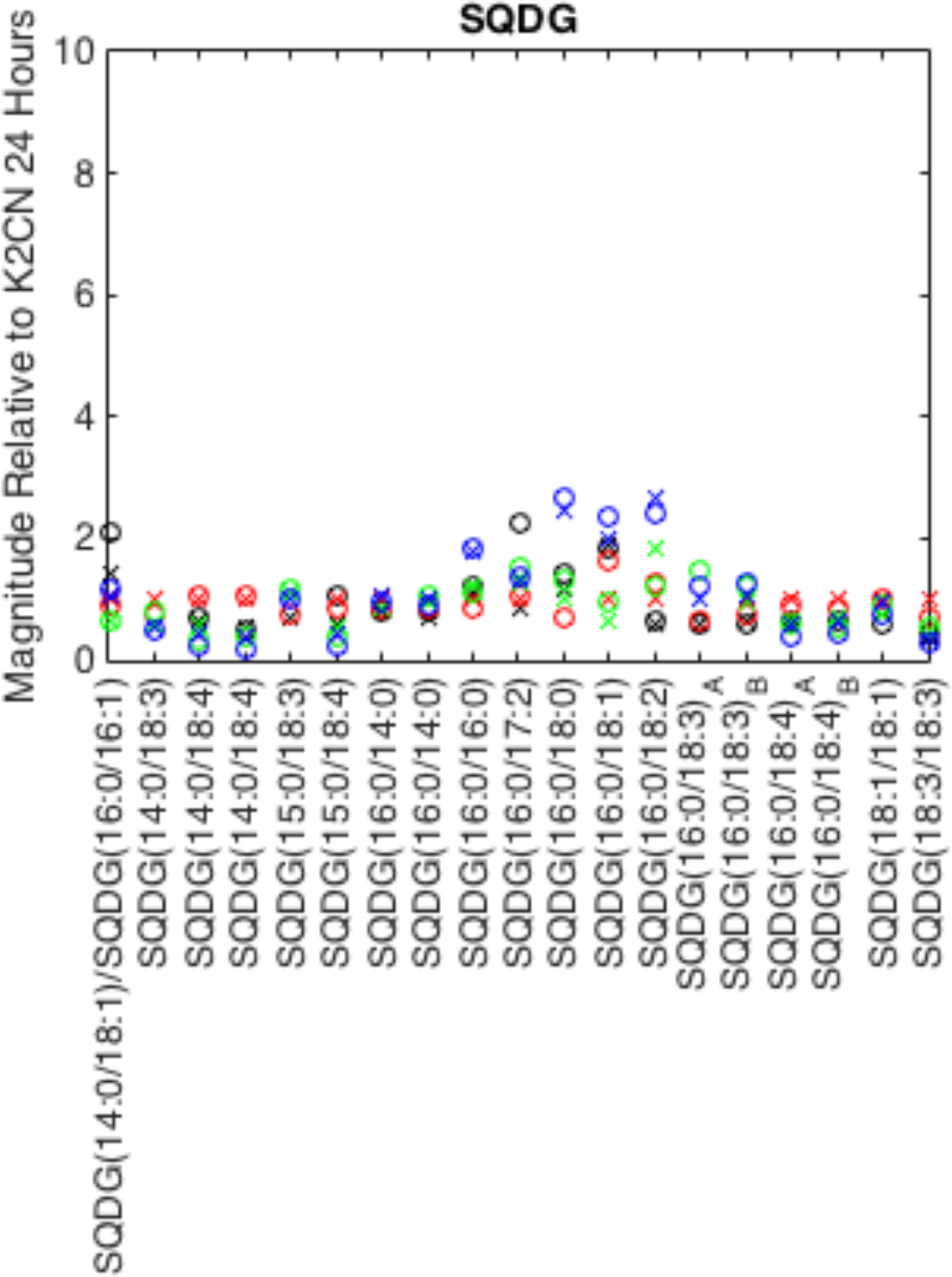
Time resolved differential expression of lipids. Sulfoquinovosyldiacylglycerol (SQDG) lipids detected via LC-MS/MS for cell cultures at 24 (X symbols) and 48 hours (O symbols) for varying C:N ratio conditions K6CN (black), K2CN (red), K2C (green) and K6C (blue) relative to K2CN at 24 hours (y-axis).

**Figure S7.**
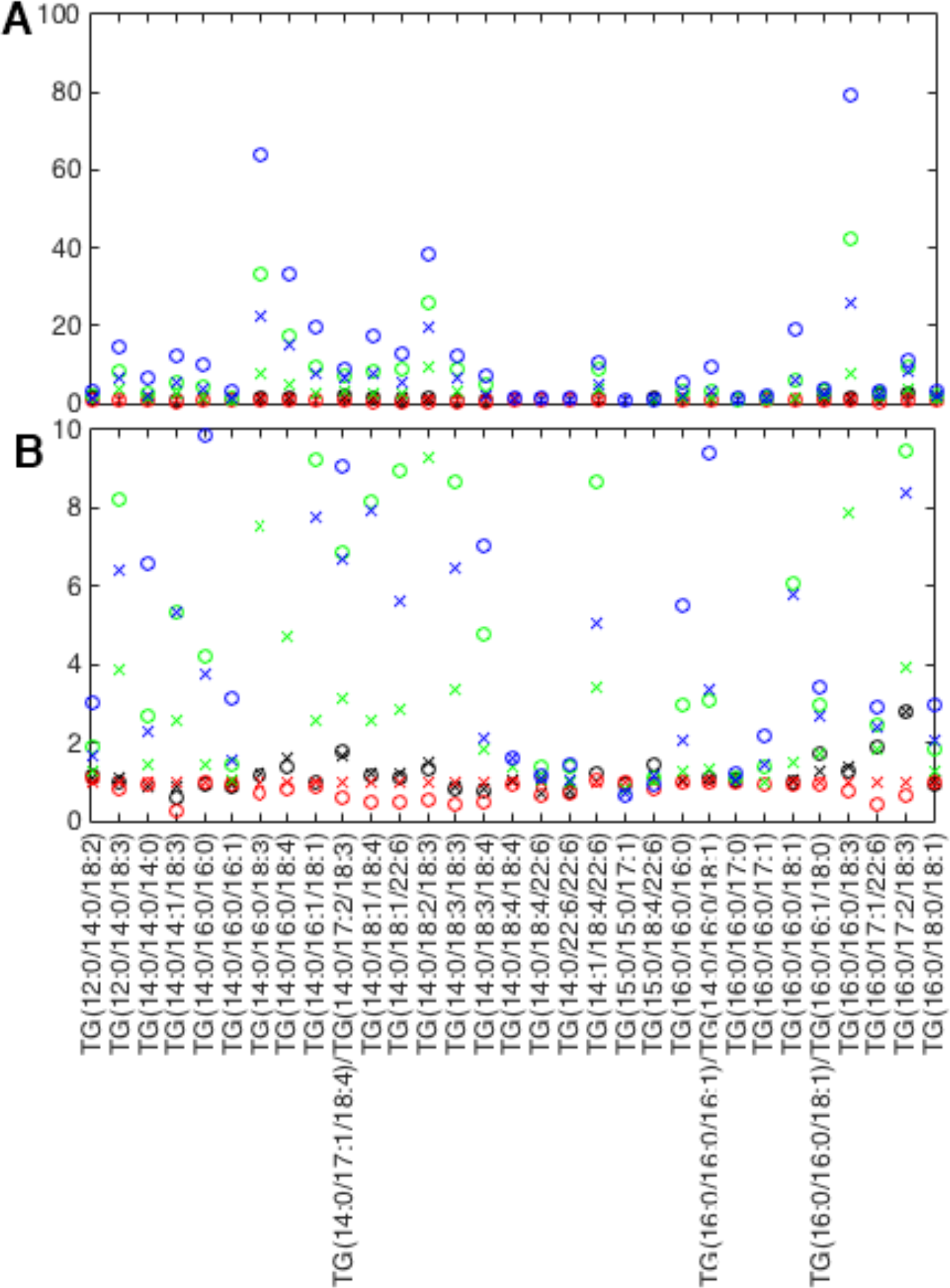
Time resolved differential abundance of triacylglycerol lipids set 1. Triacylglycerol (TG) lipidomics for cell cultures at 24 (X symbols) and 48 hours (O symbols) for varying C:N ratio conditions K6CN (black), K2CN (red), K2C (green) and K6C (blue) relative to K2CN at 24 hours (y-axis).

**Figure S8.**
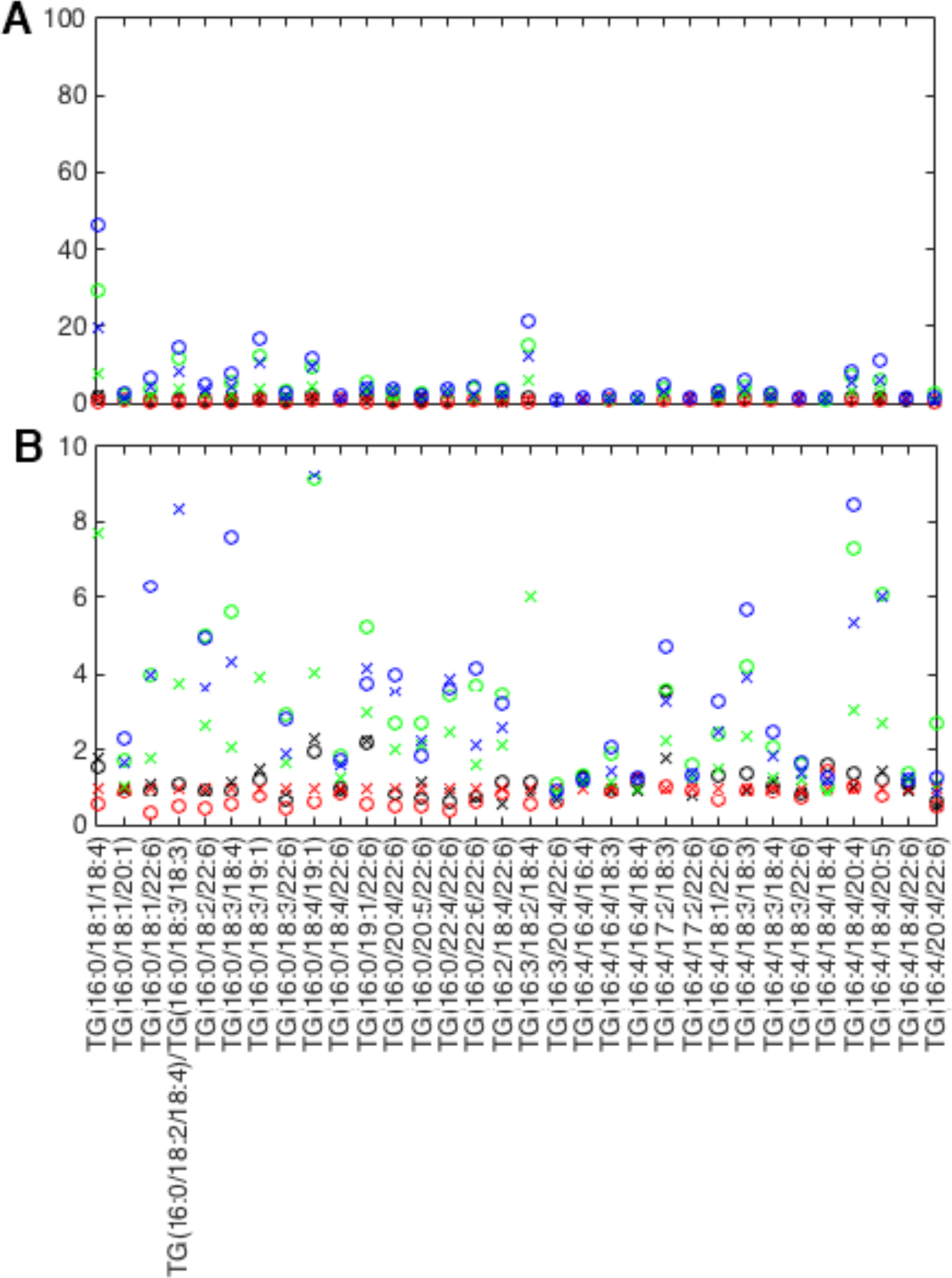
Time resolved differential abundance of triacylglycerol lipids, set 2. Triacylglycerol (TG) lipidomics detected via LC-MS/MS for cell cultures at 24 (X symbols) and 48 hours (O symbols) for varying C:N ratio conditions K6CN (black), K2CN (red), K2C (green) and K6C (blue) relative to K2CN at 24 hours (y-axis).

**Figure S9.**
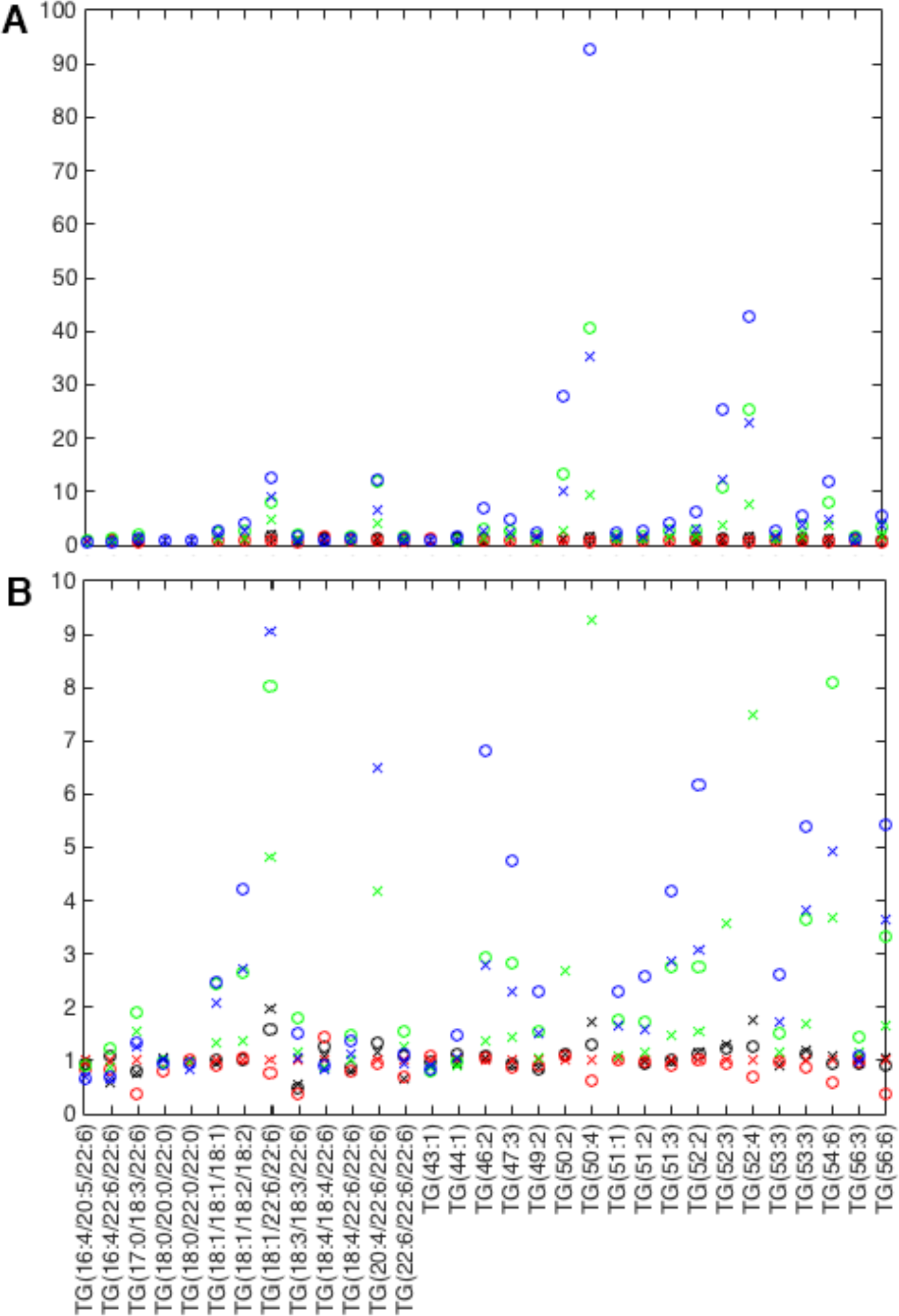
Time resolved differential abundance of long chain triacylglycerol lipids, scale 10. Long chain triacylglycerol (TG) lipidomics for cell cultures at 24 (X symbols) and 48 hours (O symbols) for varying C:N ratio conditions K6CN (black), K2CN (red), K2C (green) and K6C (blue) relative to K2CN at 24 hours (y-axis).

## References

1 Gimpel, J. A., Henriquez, V. & Mayfield, S. P. In Metabolic Engineering of Eukaryotic Microalgae: Potential and Challenges Come with Great Diversity. Front Microbiol 6, 1376, doi:10.3389/fmicb.2015.01376 (2015).

2 Wase, N., Tu, B., Allen, J. W., Black, P. N. & DiRusso, C. C. Identification and Metabolite Profiling of Chemical Activators of Lipid Accumulation in Green Algae. Plant Physiol 174, 2146–2165, doi:10.1104/pp.17.00433 (2017).

3 Longworth, J., Wu, D., Huete-Ortega, M., Wright, P. C. & Vaidyanathan, S. Proteome response of Phaeodactylum tricornutum, during lipid accumulation induced by nitrogen depletion. Algal Res 18, 213–224, doi:10.1016/j.algal.2016.06.015 (2016).

4 Levering, J., Broddrick, J. & Zengler, K. Engineering of oleaginous organisms for lipid production. Curr Opin Biotechnol 36, 32–39, doi:10.1016/j.copbio.2015.08.001 (2015).

5 Liao, J. C., Mi, L., Pontrelli, S. & Luo, S. Fuelling the future: microbial engineering for the production of sustainable biofuels. Nat Rev Microbiol 14, 288–304, doi:10.1038/nrmicro.2016.32 (2016).

6 Goncalves, E. C., Wilkie, A. C., Kirst, M. & Rathinasabapathi, B. Metabolic regulation of triacylglycerol accumulation in the green algae: identification of potential targets for engineering to improve oil yield. Plant Biotechnol J 14, 1649–1660, doi:10.1111/pbi.12523 (2016).

7 Klok, A. J., Martens, D. E., Wijffels, R. H. & Lamers, P. P. Simultaneous growth and neutral lipid accumulation in microalgae. Bioresour Technol 134, 233–243, doi:10.1016/j.biortech.2013.02.006 (2013).

8 Griffiths, M. J., van Hille, R. P. & Harrison, S. T. L. Lipid productivity, settling potential and fatty acid profile of 11 microalgal species grown under nitrogen replete and limited conditions. J Appl Phycol 24, 989–1001 (2012).

9 Guarnieri, M. T., Nag, A., Yang, S. & Pienkos, P. T. Proteomic analysis of Chlorella vulgaris: potential targets for enhanced lipid accumulation. J Proteomics 93, 245–253, doi:10.1016/j.jprot.2013.05.025 (2013).

10 Minhas, A. K., Hodgson, P., Barrow, C. J. & Adholeya, A. A Review on the Assessment of Stress Conditions for Simultaneous Production of Microalgal Lipids and Carotenoids. Front Microbiol 7, 546, doi:10.3389/fmicb.2016.00546 (2016).

11 Martin, S. F., Munagapati, V. S., Salvo-Chirnside, E., Kerr, L. E. & Le Bihan, T. Proteome turnover in the green alga Ostreococcus tauri by time course 15N metabolic labeling mass spectrometry. J Proteome Res 11, 476–486, doi:10.1021/pr2009302 (2012).

12 Lohman, E. J. et al. Optimized inorganic carbon regime for enhanced growth and lipid accumulation in Chlorella vulgaris. Biotechnol Biofuels 8, 82, doi:10.1186/s13068-015-0265-4 (2015).

13 Levering, J., Dupont, C. L., Allen, A. E., Palsson, B. O. & Zengler, K. Integrated Regulatory and Metabolic Networks of the Marine Diatom Phaeodactylum tricornutum Predict the Response to Rising CO2 Levels. mSystems 2, doi:10.1128/mSystems.00142-16 (2017).

14 Chen, Y., Xu, C. & Vaidyanathan, S. Microalgae: a robust "green bio-bridge" between energy and environment. Crit Rev Biotechnol, 1–18, doi:10.1080/07388551.2017.1355774 (2017).

15 O’Neill, J. S. et al. Circadian rhythms persist without transcription in a eukaryote. Nature 469, 554–558, doi:10.1038/nature09654 (2011).

16 Leliaert, F., Verbruggen, H. & Zechman, F. W. Into the deep: new discoveries at the base of the green plant phylogeny. Bioessays 33, 683–692, doi:10.1002/bies.201100035 (2011).

17 Courties, C. et al. Phylogenetic analysis and genome size of Ostreococcus tauri (Chlorophyta, Prasinophyceae). J Phycol 34, 844–849, doi:DOI 10.1046/j.1529-8817.1998.340844.x (1998).

18 Guillou, L. et al. Diversity of picoplanktonic prasinophytes assessed by direct nuclear SSU rDNA sequencing of environmental samples and novel isolates retrieved from oceanic and coastal marine ecosystems. Protist 155, 193–214, doi:10.1078/143446104774199592 (2004).

19 Cardol, P. et al. An original adaptation of photosynthesis in the marine green alga Ostreococcus. Proc Natl Acad Sci U S A 105, 7881–7886, doi:10.1073/pnas.0802762105 (2008).

20 Derelle, E. et al. Genome analysis of the smallest free-living eukaryote Ostreococcus tauri unveils many unique features. Proc Natl Acad Sci U S A 103, 11647–11652, doi:10.1073/pnas.0604795103 (2006).

21 Blanc-Mathieu, R. et al. Population genomics of picophytoplankton unveils novel chromosome hypervariability. Sci Adv 3, e1700239, doi:10.1126/sciadv.1700239 (2017).

22 Smallwood, C. R. et al. <em>Ostreococcus tauri</em> is a high-lipid content green algae that extrudes clustered lipid droplets. bioRxiv, doi:10.1101/249052 (2018).

23 Le Bihan, T. et al. Label-free quantitative analysis of the casein kinase 2-responsive phosphoproteome of the marine minimal model species Ostreococcus tauri. Proteomics 15, 4135–4144, doi:10.1002/pmic.201500086 (2015).

24 McCoy, J. G. et al. Discovery of sarcosine dimethylglycine methyltransferase from Galdieria sulphuraria. Proteins 74, 368–377, doi:10.1002/prot.22147 (2009).

25 Luo, G. Z., Blanco, M. A., Greer, E. L., He, C. & Shi, Y. DNA N(6)-methyladenine: a new epigenetic mark in eukaryotes? Nat Rev Mol Cell Biol 16, 705–710, doi:10.1038/nrm4076 (2015).

26 Degraeve-Guilbault, C. et al. Glycerolipid Characterization and Nutrient Deprivation-Associated Changes in the Green Picoalga Ostreococcus tauri. Plant Physiol 173, 2060–2080, doi:10.1104/pp.16.01467 (2017).

27 Shimojima, M. & Ohta, H. Critical regulation of galactolipid synthesis controls membrane differentiation and remodeling in distinct plant organs and following environmental changes. Prog Lipid Res 50, 258–266, doi:10.1016/j.plipres.2011.03.001 (2011).

28 Schaller-Laudel, S. et al. Influence of thylakoid membrane lipids on the structure of aggregated light-harvesting complexes of the diatom Thalassiosira pseudonana and the green alga Mantoniella squamata. Physiol Plant 160, 339–358, doi:10.1111/ppl.12565 (2017).

29 Losh, J. L., Young, J. N. & Morel, F. M. Rubisco is a small fraction of total protein in marine phytoplankton. The New phytologist 198, 52–58, doi:10.1111/nph.12143 (2013).

30 Noordally, Z. B. et al. Circadian protein regulation in the green lineage I. A phospho-dawn anticipates light onset before proteins peak in daytime. bioRxiv, doi:10.1101/287862 (2018).

31 Keller, M. D., Selvin, R. C., Claus, W. & Guillard, R. R. L. Media for the Culture of Oceanic Ultraphytoplankton. J Phycol 23, 633–638 (1987).

32 Ji, Y. et al. Raman spectroscopy provides a rapid, non-invasive method for quantitation of starch in live, unicellular microalgae. Biotechnol J 9, 1512–1518, doi:10.1002/biot.201400165 (2014).

33 Le Gros, M. A. et al. Biological soft X-ray tomography on beamline 2.1 at the Advanced Light Source. J Synchrotron Radiat 21, 1370–1377, doi:10.1107/S1600577514015033 (2014).

34 Parkinson, D. Y., Knoechel, C., Yang, C., Larabell, C. A. & Le Gros, M. A. Automatic alignment and reconstruction of images for soft X-ray tomography. J Struct Biol 177, 259–266, doi:10.1016/j.jsb.2011.11.027 (2012).

35 Dautel, S. E. et al. Lipidomics reveals dramatic lipid compositional changes in the maturing postnatal lung. Sci Rep 7, 40555, doi:10.1038/srep40555 (2017).

36 Kyle, J. E. et al. LIQUID: an-open source software for identifying lipids in LC-MS/MS-based lipidomics data. Bioinformatics 33, 1744–1746, doi:10.1093/bioinformatics/btx046 (2017).

37 Wang, Y. et al. Reversed-phase chromatography with multiple fraction concatenation strategy for proteome profiling of human MCF10A cells. Proteomics 11, 2019–2026, doi:10.1002/pmic.201000722 (2011).

38 Eng, J. K., McCormack, A. L. & Yates, J. R. An approach to correlate tandem mass spectral data of peptides with amino acid sequences in a protein database. J Am Soc Mass Spectrom 5, 976–989, doi:10.1016/1044-0305(94)80016-2 (1994).

39 Kim, S., Gupta, N. & Pevzner, P. A. Spectral probabilities and generating functions of tandem mass spectra: a strike against decoy databases. J Proteome Res 7, 3354–3363, doi:10.1021/pr8001244 (2008).

40 Monroe, M. E., Shaw, J. L., Daly, D. S., Adkins, J. N. & Smith, R. D. MASIC: a software program for fast quantitation and flexible visualization of chromatographic profiles from detected LC-MS(/MS) features. Comput Biol Chem 32, 215–217, doi:10.1016/j.compbiolchem.2008.02.006 (2008).

